# Pulmonary primary oxysterol and bile acid synthesis as a predictor of outcomes in pulmonary arterial hypertension

**DOI:** 10.1101/2024.01.20.576474

**Authors:** Mona Alotaibi, Lloyd D. Harvey, William C. Nichols, Michael W. Pauciulo, Anna Hemnes, Tao Long, Jeramie D. Watrous, Arjana Begzati, Jaakko Tuomilehto, Aki S. Havulinna, Teemu J. Niiranen, Pekka Jousilahti, Veikko Salomaa, Thomas Bertero, Nick H. Kim, Ankit A. Desai, Atul Malhotra, Jason X.-J. Yuan, Susan Cheng, Stephen Y. Chan, Mohit Jain

**Author notes:** Equal contribution (Stephen Y. Chan, Mohit Jain). Current address: Sapient Bioanalytics, LLC, San Diego, CA. Corresponding Authors: Mohit Jain, MD, PhD (**Contact**) Department of Medicine University of California San Diego Biomedical Research Facility II 9500 Gilman Drive La Jolla, CA, USA 92037, Stephen Y. Chan, MD, PhD, Center for Pulmonary Vascular Biology and Medicine Pittsburgh Heart, Lung, and Blood Vascular Medicine Institute Division of Cardiology, Department of Medicine University of Pittsburgh Medical Center 200 Lothrop Street, BST E1240 Pittsburgh, PA USA 15213.

## Abstract

Pulmonary arterial hypertension (PAH) is a rare and fatal vascular disease with heterogeneous clinical manifestations. To date, molecular determinants underlying the development of PAH and related outcomes remain poorly understood. Herein, we identify pulmonary primary oxysterol and bile acid synthesis (PPOBAS) as a previously unrecognized pathway central to PAH pathophysiology. Mass spectrometry analysis of 2,756 individuals across five independent studies revealed 51 distinct circulating metabolites that predicted PAH-related mortality and were enriched within the PPOBAS pathway. Across independent single-center PAH studies, PPOBAS pathway metabolites were also associated with multiple cardiopulmonary measures of PAH-specific pathophysiology. Furthermore, PPOBAS metabolites were found to be increased in human and rodent PAH lung tissue and specifically produced by pulmonary endothelial cells, consistent with pulmonary origin. Finally, a poly-metabolite risk score comprising 13 PPOBAS molecules was found to not only predict PAH-related mortality but also outperform current clinical risk scores. This work identifies PPOBAS as specifically altered within PAH and establishes needed prognostic biomarkers for guiding therapy in PAH.

**One-Sentence Summary:** This work identifies pulmonary primary oxysterol and bile acid synthesis as altered in pulmonary arterial hypertension, thus establishing a new prognostic test for this disease.

## Introduction

Pulmonary arterial hypertension (PAH) is a complex and heterogenous vascular disease with idiopathic, heritable, autoimmune, and comorbid etiologies.(*1*) Despite significant therapeutic advances over the last few decades,(*2*) PAH still carries substantial morbidity and mortality(*3*) particularly for patients with advanced disease. Once symptomatic, the average 5-year survival ranges between 27% and 57%,(*4*) comparable to the median survival of advanced cancer.(*5*) To date, however, the molecular determinants of PAH and PAH-related outcomes remain poorly understood. Human genetics and genome-wide association studies have implicated several pathways and molecules in the pathogenesis of endothelial dysfunction and PAH, including the bone morphogenetic protein receptor type II (BMPRII), transforming growth factor-β (TGF-β), potassium two pore domain channel subfamily K member 3 (KCNK3), eukaryotic translation initiation factor 2 alpha kinase 4 (EIF2AK4), T-box transcription factor 4 (TBX4), SRY-box transcription factor 17 (SOX17), major histocompatibility complex, class II, DP alpha 1 (HLA-DPA1), and nitric oxide signaling.(*6–10*) However, measurable alterations across these pathways have proven to be negligible predictors of PAH mortality. Furthermore, pharmacologic control of these pathways has led to minimal improvements in PAH survival, suggesting that additional critical molecular modulators of PAH pathobiology and prognosis may exist.

Recent attention has centered on pulmonary vascular metabolic dysregulation as a potential key driver of PAH pathogenesis and severity.(*11*) Interrogating the metabolic regulation of vascular integrity and function is now made possible by comprehensively profiling circulating small molecules that reflect homeostatic or perturbed pathway activity across human disease states (“metabolomics”).(*12–16*) Accordingly, plasma metabolomic signatures of various clinical traits and outcomes have offered insights into mechanisms of human disease, provided prognostic indicators, and served as targets and measures of response to novel therapies.(*14, 15*) Prior efforts to leverage metabolomics for the study of PAH have been limited to small-sized, single center cohorts and have focused on predominantly well-described metabolic pathways.(*17–19*) More recently, advances in bioanalytical technologies and systems biology approaches have made it possible to not only broaden the range of measurable bioactive small molecules but also facilitate the identification of novel (*i.e*., previously non-annotated) metabolites that may be identified in relation to a particular disease phenotype or outcome of interest.

Herein, we leverage state-of-the-art mass spectrometry-based metabolomics platform, applied to the largest known collection of human PAH biosamples amassed to date across multi-center discovery and validation cohorts. Through multi-dimensional integration of genomic measures with tens of thousands of annotated and non-annotated circulating metabolites, we identify a key molecular association between primary pulmonary oxysterol and bile acid synthesis (PPOBAS) and PAH pathobiology, including disease severity and overall prognosis. Furthermore, our accompanying study (by Harvey et al.)(*20*) provides a mechanistic insight and causal role for PPOBAS as inflammatory driver in the severity of PAH. This work highlights a previously unrecognized pathway underlying PAH and establishes more effective prognostic tools that can be used to guide the development of emerging therapies in human PAH.

## Results

An overview of the study design is provided in **Fig.1**. At the outset, biosamples from a total of 2,756 participants of five independent studies of human PAH and non-PAH controls were assembled with demographic and clinical characteristics including PAH-relevant cardiopulmonary hemodynamic measures summarized in **Table 1**. All PAH patients were under surveillance for clinical outcomes including mortality; of the 2,712 patients with PAH enrolled at baseline, a total of 578 deaths (21.3% mortality rate) occurred during the study period.

**Fig. 1.**
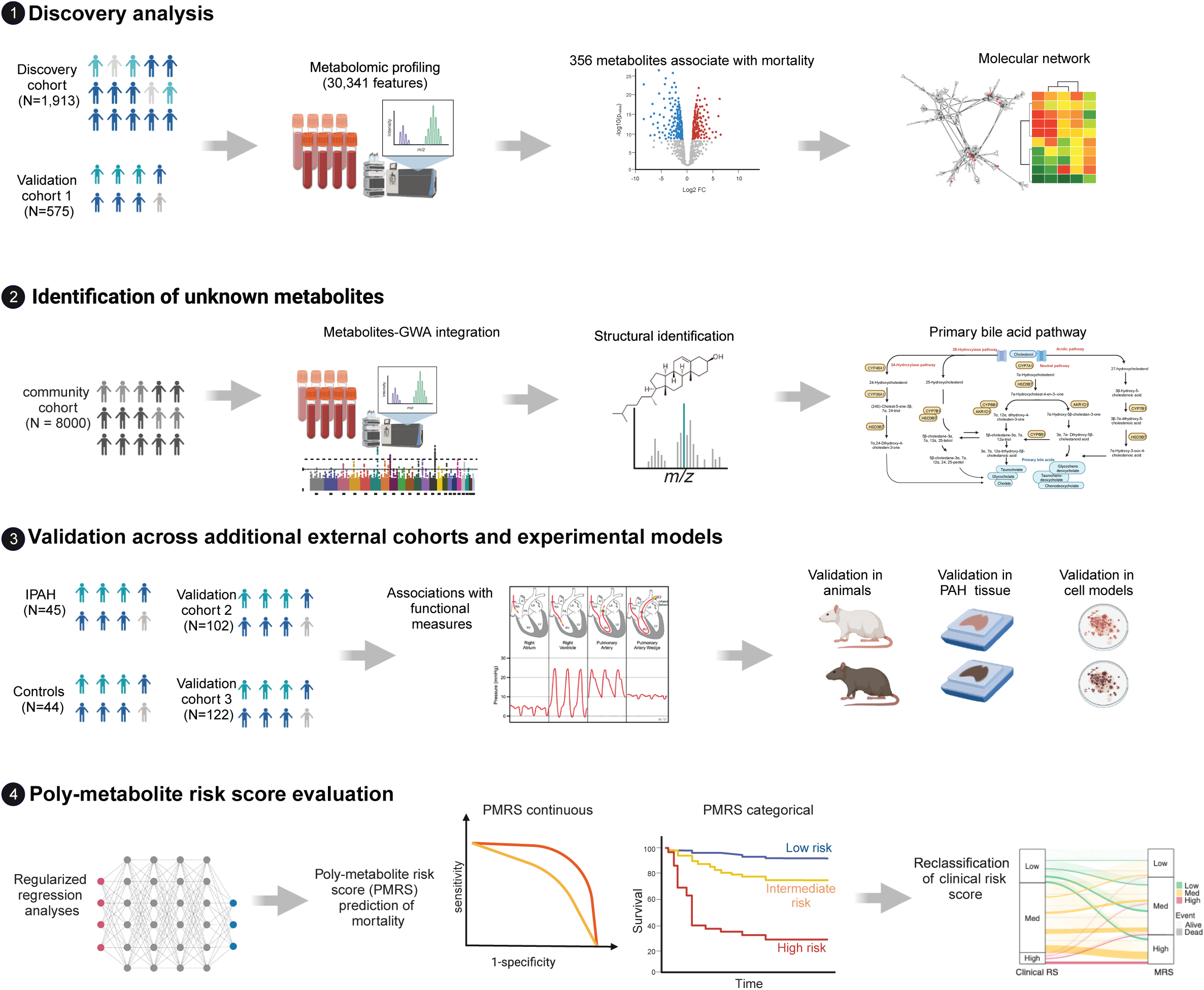
Overall study design.

**Table 1.**
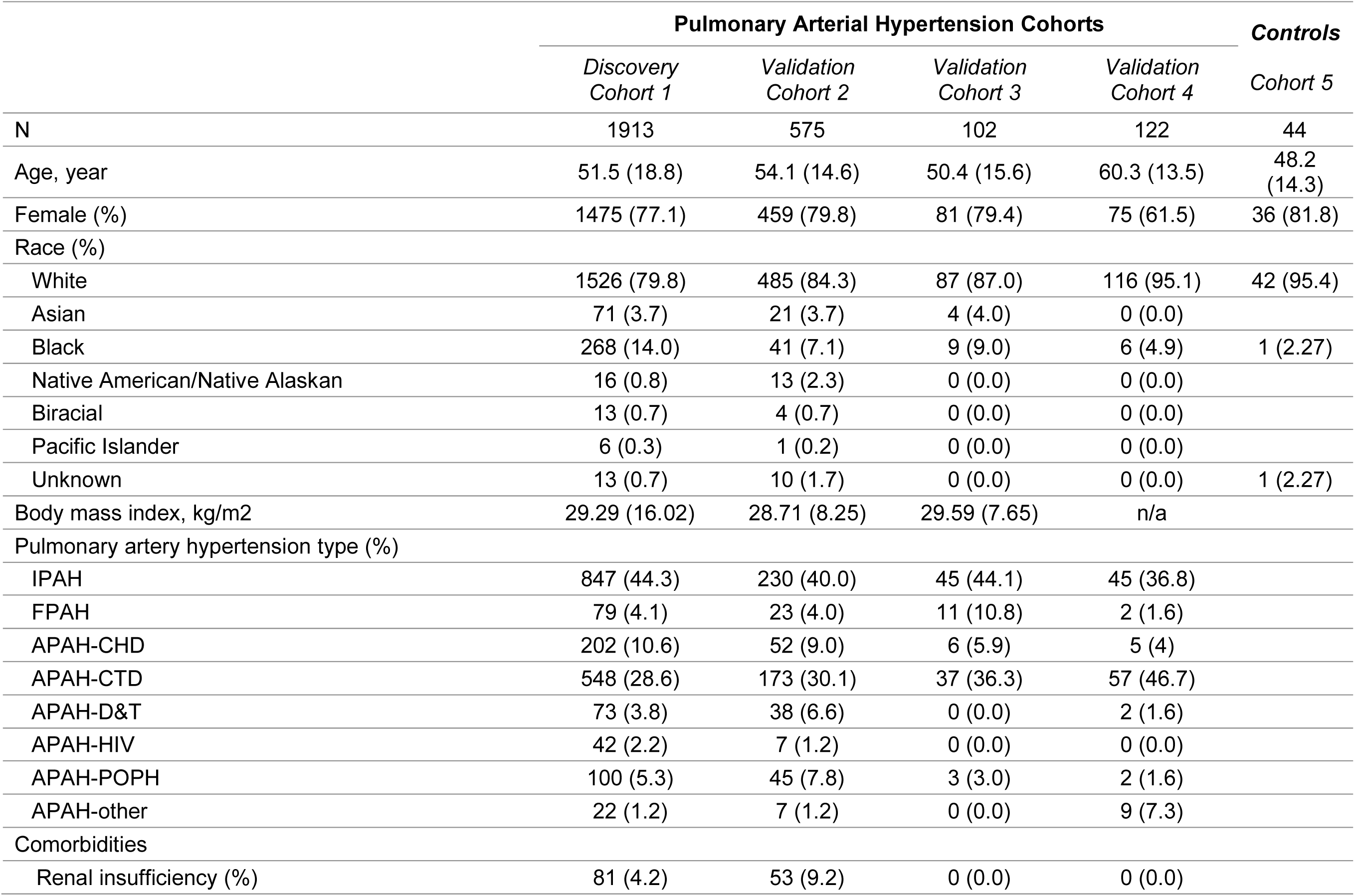
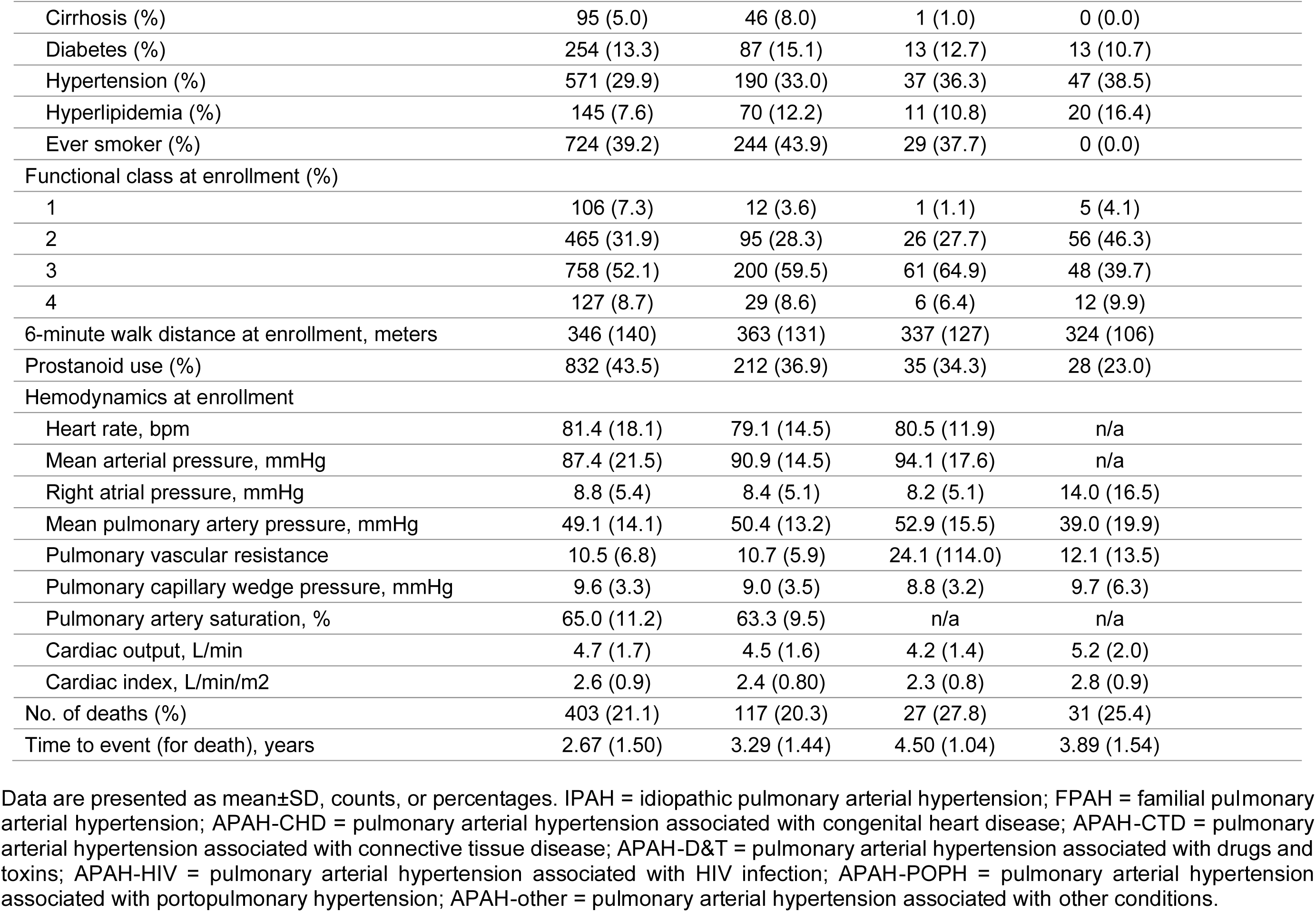
Characteristics of the PAH discovery and validation cohorts.

|  | **Pulmonary Arterial Hypertension Cohorts** |  |  |  | **Controls** |
| --- | --- | --- | --- | --- | --- |
|  | *Discovery Cohort 1* | *Validation Cohort 2* | *Validation Cohort 3* | *Validation Cohort 4* | *Cohort 5* |
| N | 1913 | 575 | 102 | 122 | 44 |
| Age, year | 51.5 (18.8) | 54.1 (14.6) | 50.4 (15.6) | 60.3 (13.5) | 48.2 (14.3) |
| Female (%) | 1475 (77.1) | 459 (79.8) | 81 (79.4) | 75 (61.5) | 36 (81.8) |
| Race (%) |  |  |  |  |  |
| White | 1526 (79.8) | 485 (84.3) | 87 (87.0) | 116 (95.1) | 42 (95.4) |
| Asian | 71 (3.7) | 21 (3.7) | 4 (4.0) | 0 (0.0) |  |
| Black | 268 (14.0) | 41 (7.1) | 9 (9.0) | 6 (4.9) | 1 (2.27) |
| Native American/Native Alaskan | 16 (0.8) | 13 (2.3) | 0 (0.0) | 0 (0.0) |  |
| Biracial | 13 (0.7) | 4 (0.7) | 0 (0.0) | 0 (0.0) |  |
| Pacific Islander | 6 (0.3) | 1 (0.2) | 0 (0.0) | 0 (0.0) |  |
| Unknown | 13 (0.7) | 10 (1.7) | 0 (0.0) | 0 (0.0) | 1 (2.27) |
| Body mass index, kg/m2 | 29.29 (16.02) | 28.71 (8.25) | 29.59 (7.65) | n/a |  |
| Pulmonary artery hypertension type (%) |  |  |  |  |  |
| IPAH | 847 (44.3) | 230 (40.0) | 45 (44.1) | 45 (36.8) |  |
| FPAH | 79 (4.1) | 23 (4.0) | 11 (10.8) | 2 (1.6) |  |
| APAH-CHD | 202 (10.6) | 52 (9.0) | 6 (5.9) | 5 (4) |  |
| APAH-CTD | 548 (28.6) | 173 (30.1) | 37 (36.3) | 57 (46.7) |  |
| APAH-D&T | 73 (3.8) | 38 (6.6) | 0 (0.0) | 2 (1.6) |  |
| APAH-HIV | 42 (2.2) | 7 (1.2) | 0 (0.0) | 0 (0.0) |  |
| APAH-POPH | 100 (5.3) | 45 (7.8) | 3 (3.0) | 2 (1.6) |  |
| APAH-other | 22 (1.2) | 7 (1.2) | 0 (0.0) | 9 (7.3) |  |
| Comorbidities |  |  |  |  |  |
| Renal insufficiency (%) | 81 (4.2) | 53 (9.2) | 0 (0.0) | 0 (0.0) |  |

|  |  |  |  |  |
| --- | --- | --- | --- | --- |
| Cirrhosis (%) | 95 (5.0) | 46 (8.0) | 1 (1.0) | 0 (0.0) |
| Diabetes (%) | 254 (13.3) | 87 (15.1) | 13 (12.7) | 13 (10.7) |
| Hypertension (%) | 571 (29.9) | 190 (33.0) | 37 (36.3) | 47 (38.5) |
| Hyperlipidemia (%) | 145 (7.6) | 70 (12.2) | 11 (10.8) | 20 (16.4) |
| Ever smoker (%) | 724 (39.2) | 244 (43.9) | 29 (37.7) | 0 (0.0) |
| Functional class at enrollment (%) |  |  |  |  |
| 1 | 106 (7.3) | 12 (3.6) | 1 (1.1) | 5 (4.1) |
| 2 | 465 (31.9) | 95 (28.3) | 26 (27.7) | 56 (46.3) |
| 3 | 758 (52.1) | 200 (59.5) | 61 (64.9) | 48 (39.7) |
| 4 | 127 (8.7) | 29 (8.6) | 6 (6.4) | 12 (9.9) |
| 6-minute walk distance at enrollment, meters | 346 (140) | 363 (131) | 337 (127) | 324 (106) |
| Prostanoid use (%) | 832 (43.5) | 212 (36.9) | 35 (34.3) | 28 (23.0) |
| Hemodynamics at enrollment |  |  |  |  |
| Heart rate, bpm | 81.4 (18.1) | 79.1 (14.5) | 80.5 (11.9) | n/a |
| Mean arterial pressure, mmHg | 87.4 (21.5) | 90.9 (14.5) | 94.1 (17.6) | n/a |
| Right atrial pressure, mmHg | 8.8 (5.4) | 8.4 (5.1) | 8.2 (5.1) | 14.0 (16.5) |
| Mean pulmonary artery pressure, mmHg | 49.1 (14.1) | 50.4 (13.2) | 52.9 (15.5) | 39.0 (19.9) |
| Pulmonary vascular resistance | 10.5 (6.8) | 10.7 (5.9) | 24.1 (114.0) | 12.1 (13.5) |
| Pulmonary capillary wedge pressure, mmHg | 9.6 (3.3) | 9.0 (3.5) | 8.8 (3.2) | 9.7 (6.3) |
| Pulmonary artery saturation, % | 65.0 (11.2) | 63.3 (9.5) | n/a | n/a |
| Cardiac output, L/min | 4.7 (1.7) | 4.5 (1.6) | 4.2 (1.4) | 5.2 (2.0) |
| Cardiac index, L/min/m<sup>2</sup> | 2.6 (0.9) | 2.4 (0.80) | 2.3 (0.8) | 2.8 (0.9) |
| No. of deaths (%) | 403 (21.1) | 117 (20.3) | 27 (27.8) | 31 (25.4) |
| Time to event (for death), years | 2.67 (1.50) | 3.29 (1.44) | 4.50 (1.04) | 3.89 (1.54) |

Data are presented as mean±SD, counts, or percentages. IPAH = idiopathic pulmonary arterial hypertension; FPAH = familial pulmonary arterial hypertension; APAH-CHD = pulmonary arterial hypertension associated with congenital heart disease; APAH-CTD = pulmonary arterial hypertension associated with connective tissue disease; APAH-D&T = pulmonary arterial hypertension associated with drugs and toxins; APAH-HIV = pulmonary arterial hypertension associated with HIV infection; APAH-POPH = pulmonary arterial hypertension associated with portopulmonary hypertension; APAH-other = pulmonary arterial hypertension associated with other conditions.

### Metabolome-wide association of plasma metabolites with PAH mortality

Using state-of-the-art, nontargeted (“discovery”) mass spectrometry, a total of 30,341 metabolite features were assayed in the discovery cohort that comprised 1,913 individuals enrolled in the PAH Biobank (www.pahbiobank.org) (**Fig. 2A**).(*10, 17*) From these comprehensive measures of circulating small molecule biomarkers, 356 metabolites demonstrated associations at a metabolome-wide level with the composite outcome of all-cause mortality or heart-lung transplantation in the discovery cohort after adjustment for age, sex, body mass index, cirrhosis, renal failure, prostacyclin therapy, aspirin, and statin therapy (Discovery *P*<1×10^-6^, **Fig. 2A** and **table S1**). Of these, 271 circulating markers were positively associated with clinical outcomes (HR 1.12 to 1.77 per 1-SD log-analyte) and confirmed in 575 individuals from within the PAH Biobank but collected at independent sites (Validation *P*<1×10^-4^, **table S1**). Moreover, this association with mortality was validated in a third independent, single center cohort of PAH patients (**table S2**). Among the 356 metabolites, we identified nervonic acid and lignoceric acid, which we have previously reported to be altered in scleroderma-associated PAH and associated with PAH disease severity.(*19*) In addition, two primary bile acids, chenodeoxycholic acid and glycocholic acid, were also found to be elevated in association with PAH mortality. The remaining molecules represented novel and not previously annotated metabolites.

**Fig. 2.**
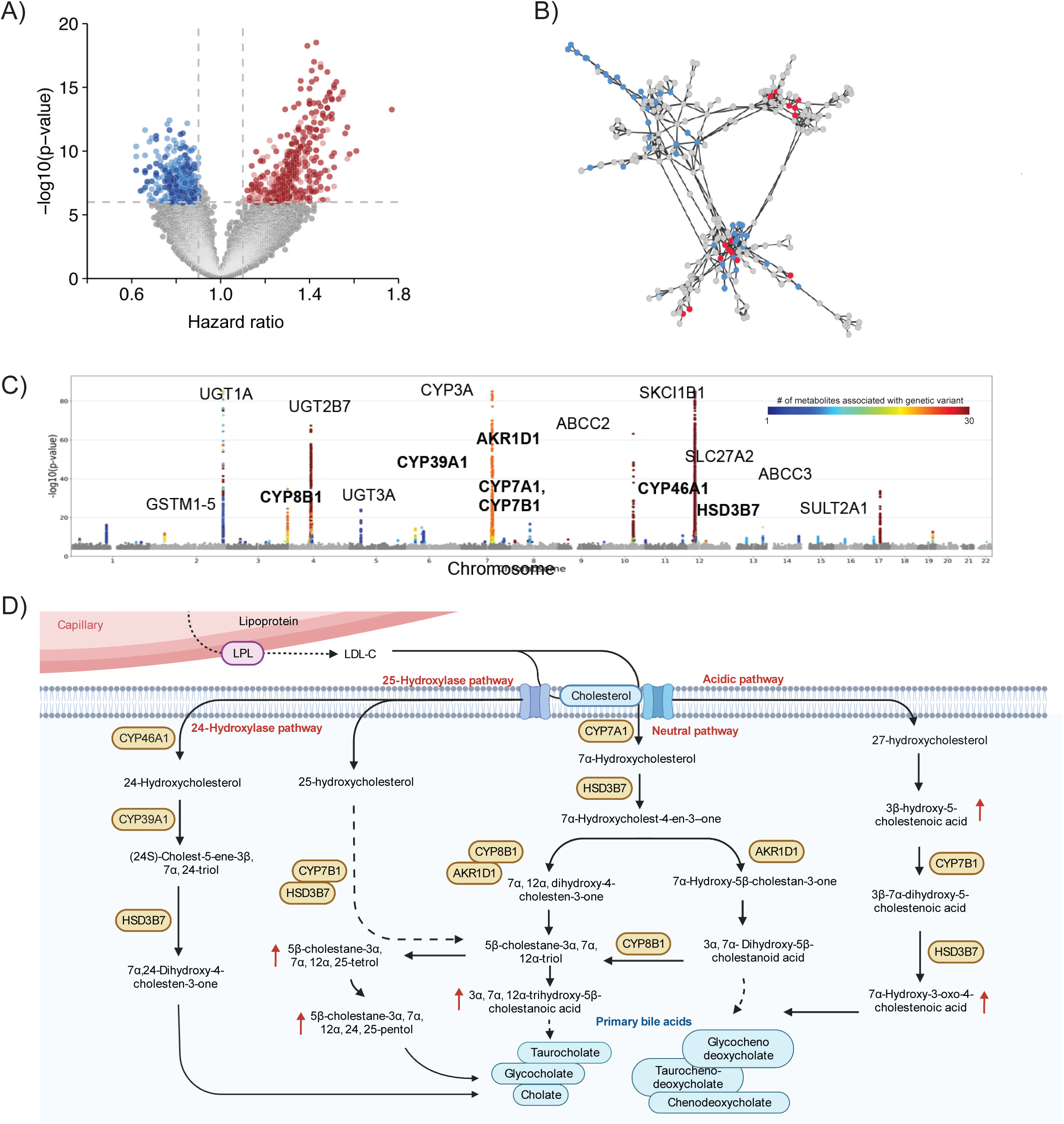
Metabolites associated with mortality in PAH discovery and validation cohorts. (**A**) Volcano plot of metabolites associated with mortality in discovery and validation cohorts. Data are derived from multivariable-adjusted Cox regression models. Red dots indicate deleterious metabolites significant in both discovery and validation cohorts; pink dots indicate deleterious metabolites significant only in the discovery cohort; dark blue dots indicate protective metabolites in both discovery and validation cohorts; light blue dots indicate protective metabolites significant only in discovery cohort; grey dots indicate all metabolites measured in discovery and validation cohorts. (**B**) Chemical networking of significant metabolites in both discovery and validation cohorts. Tandem mass spectra for the chromatographic features were clustered against tandem mass spectra for commercial standards with the resulting correlations displayed as a nested molecular network. Here, red nodes indicate tandem mass spectra for commercial standards, blue nodes indicate tandem mass spectra from glucuronidated compounds and grey nodes indicate tandem mass spectra from the 356 unknown compounds. (**C**) Manhattan plot of associations between genetic variants and glucuronidated 51 metabolite features. For single nucleotide polymorphisms (SNPs) with multiple associations, only the lowest *P* value is shown. The y-axis is truncated at −log10(*P*) for improved visualization, and SNPs with *P*>10^−7^ are omitted. Color scale reflect the number of metabolites associated with genetic variant. Top scale (dark red) indicates 30 metabolite and lower scale (dark blue) indicates one metabolite. (**D**) Schematic representation of primary bile acid synthesis pathway. Key enzymes associated with the glucuronidated 51 metabolite features are highlighted in yellow.

### Identification of novel metabolites observed in association with PAH mortality

To identify the biological pathways enriched among the metabolites found in association with PAH mortality, which comprised predominantly novel (previously non-annotated) molecules, spectral network analysis was performed with clustering of metabolites according to MS/MS chemical fingerprints (**Fig. 2B**).(*21–24*) Approximately 15% of all PAH mortality-associated metabolites (51/356) demonstrated close chemical similarity with a shared sub-fragment identified as a common glucuronide modification (**fig. S1**), suggesting a common biological process central to PAH. To identify the biologic origins and relationships among the novel unknown metabolites, we employed genomic integration with genome-wide association of metabolites. Across an independent study of ∼8,000 community-dwelling adults, from the FINRISK-2002 study, with extensive genotype data in addition to detailed assays of circulating metabolite levels,(*25, 26*) 30 of the 51 prioritized metabolites were found to correspond with genetic variants in haplotypes with or adjacent to genes encoding key enzymes within the primary oxysterol and bile acid synthesis pathway, including *AKR1D1*, *CYP7A1*, *CYP7B1*, *CYP8B1*, *CYP39A1*, *CYP46A1*, *HSD3B7*, and *SLC27A2*, as well as with enzymes required for glucuronidation and transport of intermediary organic sterols, including *UGT1A6*, *UGT2A*, *UGT2B*, *ABCC*, *SLCO1A2,* and *SC5D* at a genome-wide significance threshold (*P*<1×10^-8^, **Fig. 2C**). Importantly, this biochemical pathway is known to be critical for primary conversion of cholesterol to oxysterol intermediates and primary bile acids, suggesting that the PAH metabolites of interest may represent glucuronidated primary bile acid and oxysterol intermediates. To elucidate the structural identity of metabolites, commercial standards for oxysterols and bile alcohol intermediates were obtained (**table S3**) and conjugated using microsomal glucuronidation.(*27*) Five of the metabolites of interest were definitively identified as glucuronidated forms of primary bile acid synthesis intermediates, including 3β-hydroxy-5-cholestanoic acid, 7α-hydroxy-3-oxo-4-cholestenoic (7-HOCA), 3α,7α,12α-trihydroxy-5β-cholestanoic acid, 5β-cholestane-3α,7α,12α,26-tetrol, and 5β-cholestane-3α,7α,12α,25,26-pentol based on precise matching of accurate mass features, chromatographic retention, and peak shapes with standards. All five of these molecules were found to be elevated in association with adverse PAH outcomes (**Fig. 2D**). Similarly, MS/MS fragmentation analysis of the remaining associated metabolites revealed 46 of the 51 molecules were consistent with putative primary bile acid metabolites and intermediates, with a high degree of correlation among all 51 molecules in patients with PAH (**fig. S2**). Collectively, these data define a signature of elevated circulating primary bile acid and oxysterol metabolites in patients with PAH and in relation to PAH-related mortality.

### Validation in external cohorts: primary oxysterol and bile acid metabolites are associated with clinical disease severity in PAH

To determine whether primary bile acid metabolites are consistently altered in relation to PAH and PAH-related physiology, oxysterols and bile acid intermediates were examined in an independent, single center study of PAH versus controls (**Table 1**). Matched control samples were obtained from both healthy volunteers as well as ‘disease controls’ presenting with symptomatic dyspnea and exercise intolerance, but not PAH confirmed by right heart catheterization. Of the 51 primary bile acid metabolites, 33 were altered with consistent directionality in individuals with PAH relative to healthy counterparts (**Fig. 3A** and **table S4**). Elevation of circulating primary bile acid metabolites was associated with key clinical measures of PAH severity and hemodynamics, including decreased six minute walk distance (6MWD, odds ratio [OR] –34 to –16, *P*<1×10^-6^), higher World Health Organization (WHO) functional class (WHO FC, OR 1.2 to 1.6, *P*<0.05), elevated mean right atrial pressure (mRAP, OR 0.05 to 0.11, *P* < 1 x 10^-4^), reduced cardiac index (CI, OR –0.3x to –0.1x, *P*<0.01), and elevated pulmonary vascular resistance (PVR, OR 0.02 to 0.05, *P*<0.05). These associations were cross-validated across independent cohorts (**Fig. 3B** and **table S5**). Taken together, circulating metabolites within the primary bile acid pathway were found to reliably predict PAH disease severity and clinical outcomes.

**Fig. 3.**
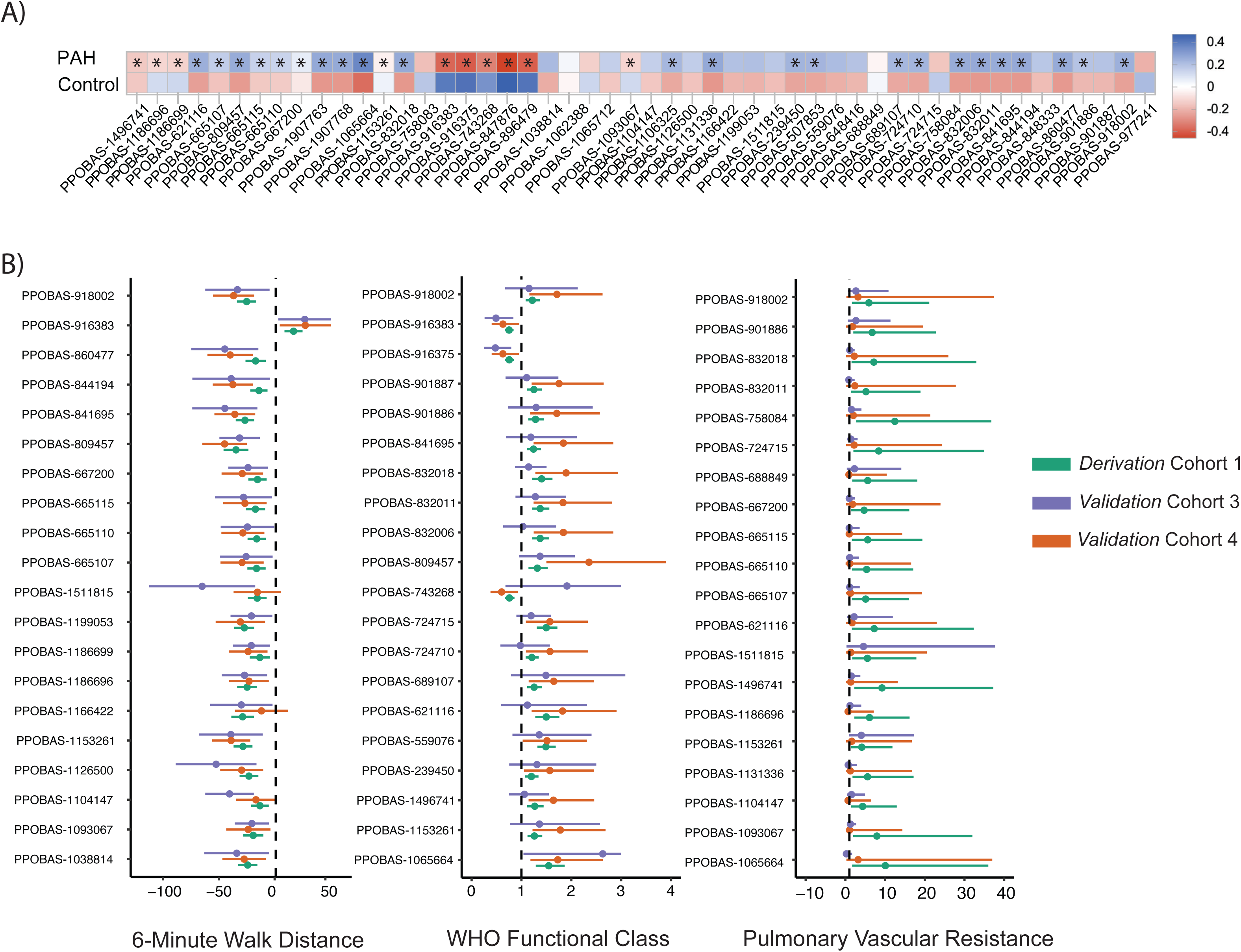
Primary bile acid metabolites associate with PAH outcomes and physiology. (**A**) Heatmap of primary bile acid and oxysterol metabolites in idiopathic PAH patients (N=45) and control plasma (N=44). Each square indicates relative abundance of metabolite value in PAH versus control. Data originated from multivariable logistic regression analysis. Asterisks (*) indicate significance with *P*<0.05. (**B**) Forest plots of the top 20 glucuronide metabolites and their association to 6-minute walk distance (meters), WHO functional class, and pulmonary vascular resistance (Wood units) in three independent cohorts of PAH (Cohort 1, N=1913; Cohort 3, N=102; Cohort 4, N=122). Data are derived from multivariable regression models adjusting for age, sex, and body mass index. All data are presented as odds ratio of clinical/hemodynamic metric per 1-SD increase in metabolite concentration.

### Validation in experimental models: dysregulated bile acid metabolite synthesis is localized to the lung endothelium in PAH

To determine whether modulation of primary bile acid metabolites is conserved in multiple established preclinical models of PAH, molecules were assayed in plasma from the SU5416 and chronic hypoxia-exposed PAH rat model (SU/Hyp)(*28*) and the monocrotaline (MCT) PAH rat model.(*29*) With the development of PAH, six of primary bile acid synthesis metabolites were found to be significantly altered in at least one of these preclinical systems (**Figs. 4A–4F**), indicating conservation of this biological process.

**Fig. 4.**
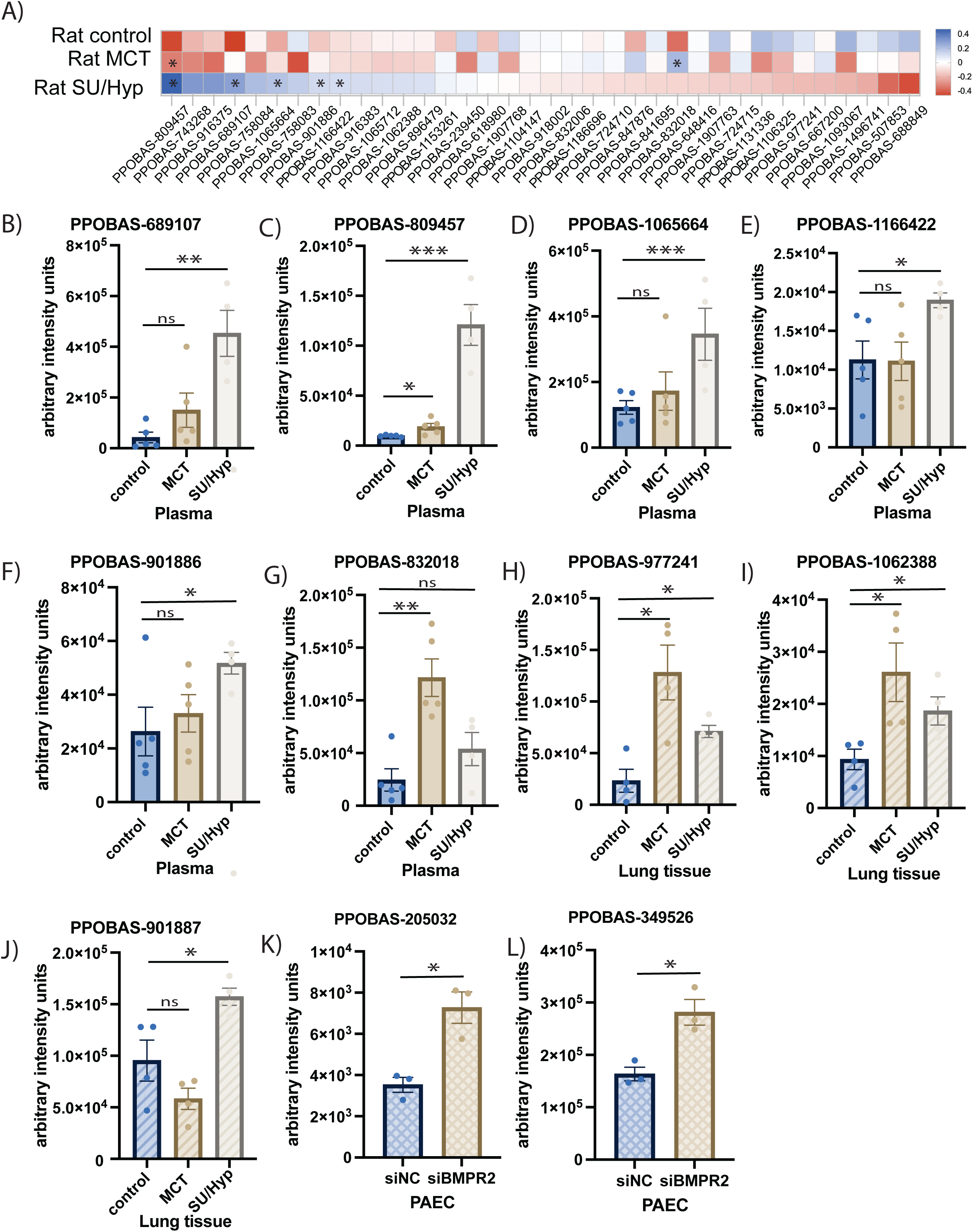
Elevation of primary oxysterol and bile acid metabolites in preclinical models of PAH. (**A**) Heatmap of oxysterol and primary bile acid metabolites in plasma samples of vehicle control, SU5146/hypoxic (SU/Hyp), or monocrotaline (MCT) PAH rat models. Each square indicates relative abundance of metabolite value. Data originated from student *t*-test. \**P*<0.05. (**B** to **G**) Significantly elevated oxysterols and primary bile acid metabolites in plasma samples of SU/Hyp or MCT rats compared to vehicle controls. (**H** to **J**) Significantly elevated oxysterols and primary bile acid metabolites in lung tissue of SU/Hyp or MCT rats compared to vehicle controls. \**P*<0.05; \*\**P*<0.01; \*\*\**P*<0.001; ns, not significant with *P*>0.05. (**K** and **L**) Significantly elevated oxysterols and primary bile acid metabolites in pulmonary artery endothelial cells. PAEC were transfected with siRNA targeting BMPR2 (siBMPR2) and compared with scrambled siRNA (siNC, n=3). Data are presented as mean±SEM. Samples were tested in triplicate. Data were analyzed with an unpaired t test. *P<0.05

Primary bile acid metabolism largely occurs within the liver, though the enzymes involved in this pathway are expressed throughput the body.(*30*) To determine whether the primary bile acid metabolites observed in PAH may be generated within the lung, tissue was isolated from the preclinical SU/Hyp, and MCT preclinical models. Across both preclinical models, primary bile acid metabolites were found to be elevated within the lung tissue (**Figs. 4H–4J**). Similarly, in explanted tissues from PAH patients and non-PAH control patients undergoing lung and heart transplantation, primary bile acid intermediates were significantly elevated in lung tissues of PAH compared to controls (**table S6**). To further determine whether bile acid metabolites may originate within the vascular endothelium, human pulmonary artery endothelial cells (PAECs) were transfected with siRNA targeting BMPR2 (bone morphogenetic protein type 2 receptor)—a major mutation in hereditary and sporadic PAH.(*6, 7*) Relative to control, knockdown of BMPR2 in PAECs resulted in higher steady-state levels of primary bile acid metabolites (**Figs. 4K–4L**). These results demonstrate that primary PPOBAS metabolites can be generated within the endothelium under the context of PAH triggers.

### A prognostic poly-metabolite risk score to predict PAH disease severity and mortality

Identification of PAH patients at greatest risk for early mortality remains clinically challenging but a major priority.(*31*) To determine the utility of PPOBAS metabolites for clinical prognostication in PAH, a poly-metabolite risk score (PAH-PMRS) was established and examined in comparison to established clinical risk metrics. Regularized survival analysis (Lasso feature selection) was used to establish a PAH-PMRS consisting of 13 PPOBAS metabolites (**table S7,** see Methods). PAH-PMRS significantly predicted PAH related mortality (AUC 73%, 95% CI [68%-78%]) with a concordance index (C-index) for 4-year all-cause mortality of 0.76 in the initial discovery study (**Table 1**), as well as in an independent external study with AUC 70% (95% CI [58%-82%]) (cohort 3) (**Fig. 5A**). By comparison, a comprehensive clinical risk score composed of common demographics, clinical metrics, and cardiac hemodynamics used in PAH risk assessment (including age, sex, body mass index, pulmonary vascular resistance, World Health Organization functional class, 6-minute walk distance, and mean right atrial pressure)(*32–36*) exhibited less predictive value with AUC of 65% (95% CI [56%-75%]). Furthermore, the conventional clinical risk score offered minimal additional predictive value to the PAH-PMRS (**Fig. 5A**). We next used the PAH-PMRS to categorize patients as low-risk (lower tertile, N=674), intermediate-risk (middle tertile, N=1180), or high-risk (top tertile, N=617). Kaplan-Meier (KM) survival analysis revealed that patients in the highest risk PAH-PMRS tertile experienced a lower overall survival rate, with 2-year survival estimated at 65% and 4-year survival estimated at 50%. The log-rank test showed a significant difference in overall survival between the risk groups (*P*<10^-5^), with a hazard ratio for the high-risk group of over 10 (10.38 high-risk vs low-risk group; 95% CI [7.26-15.22]) (**Figs. 5B** and **5C**). We next evaluated the reclassification ability of the PAH-PMRS compared to the European Society of Cardiology/European Respiratory Society (ESC/ERS) clinical risk score. In patients who died and were initially classified as either low– or intermediate-risk by ESC/ERS, the PAH-PMRS correctly re-stratified 38% of identified low-risk and 43% of intermediate-risk into the highest-risk PAH-PMRS tertile, providing further consistent evidence of greater predictive value offered by the PAH-PMRS (**Figs. 5D** and **5E**). Accordingly, the PAH-PMRS allowed for similar degrees of reclassification in additional independent single center study cohorts (**fig. S3**). Collectively, these results suggest that PPOBAS markers may be leveraged for risk stratification and clinical prognostication among patients with PAH.

**Fig. 5.**
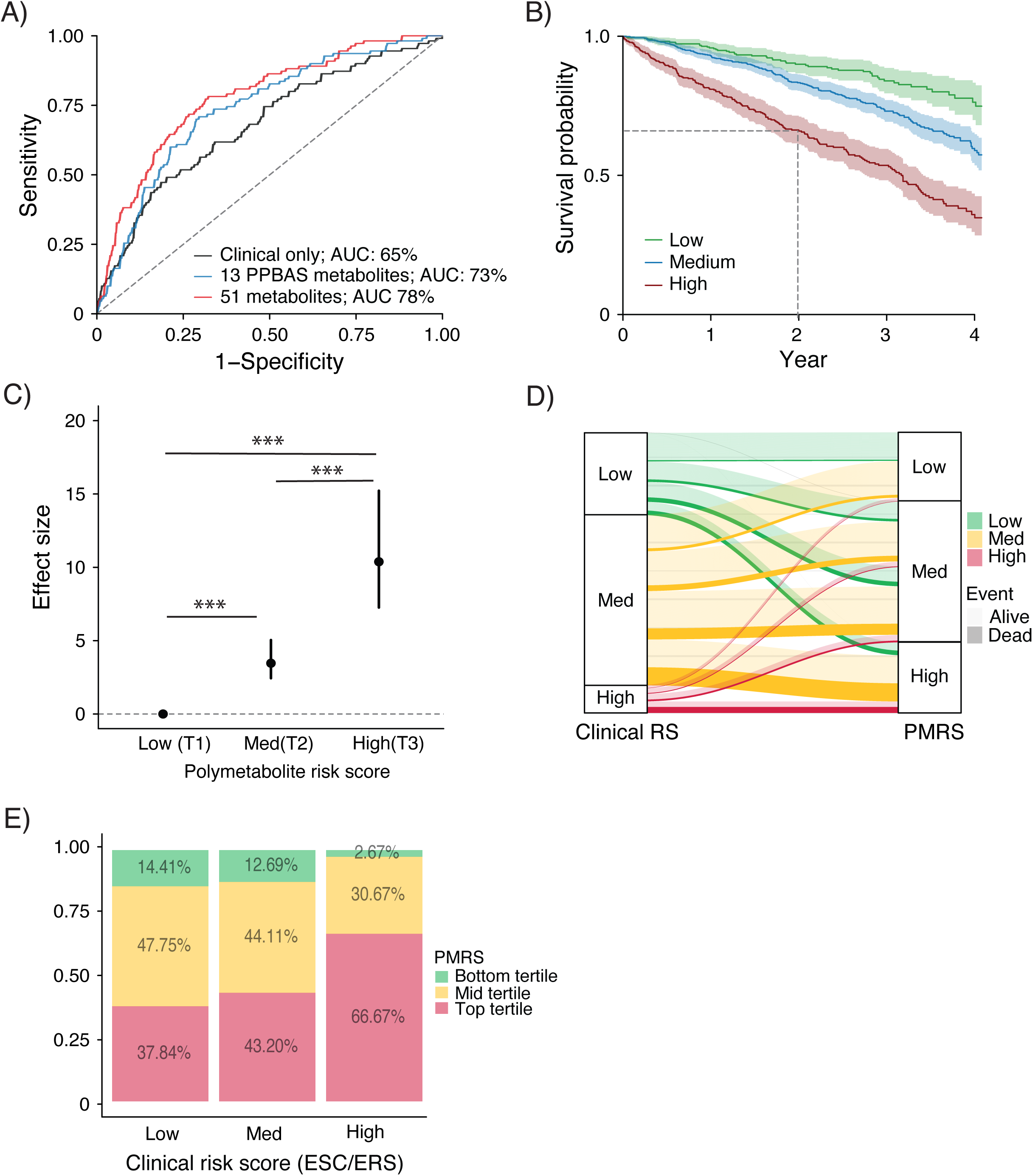
A poly-metabolite risk score predicts PAH disease severity and mortality. **(A)** Receiver-operating characteristic curves display the performance of models predicting mortality: the clinical model alone is represented by the black curve, the 13 PPOBAS poly-metabolite model by the blue curve, and 51 PPOBAS poly-metabolite model by the red curve. (**B**) Kaplan-Meier analyses display survival over time in patients with PAH within the discovery (Cohort 1) and validation (Cohort 2) cohorts, classified according to the PAH polymetabolite risk score (PAH-PMRS). (**C**) Difference in overall survival between PAH-PMRS groups normalized to low-risk group. T indicates tertile with T1 representing lower tertile; T2, middle tertile; and T3, top tertile. *P* values originated from log rank test. \*\*\**P*<0.001. (**D**) Reclassification of PAH clinical risk score (based on the European Society of Cardiology/European Respiratory Society [ESC/ERS] PAH risk calculator) using the PAH poly-metabolite risk score (PAH-PMRS). (**E**) Reclassification of PAH clinical risk score (European Society of Cardiology/European Respiratory Society [ESC/ERS]) in patients who died (N=520) in discovery (Cohort 1) and validation (Cohort 2) cohorts using the PAH poly-metabolite risk score (PAH-PMRS).

## Discussion

Leveraging a state-of-the-art mass spectrometry and an unbiased, integrative, multi-omics approach, we identified new molecular pathways associated with PAH. Representing the most comprehensive metabolomic analysis of PAH clinical samples to date, our analyses identified 51 distinct PPOBAS pathway metabolites in association with a composite outcome of all-cause mortality or heart-lung transplantation, as well as with measures of clinical disease severity and cardiopulmonary physiology across discovery and multiple independent validation cohorts. Furthermore, we found conservation of the PPOBAS associations in preclinical models of PAH with PPOBAS metabolites found to be elevated in lung tissue and specifically produced by pulmonary vascular endothelium *in vitro*, suggesting a potential role in disease pathobiology. Finally, we generated a prognostic PAH-PMRS that outperformed conventional metrics for prognostication of PAH across independent clinical cohorts. Our accompanying study (by Harvey et al.)(*20*) elucidates a causal role of PPOBAS within the endothelium in the progression and severity of PAH. Together, these findings identify PPOBAS as a previously unrecognized biological pathway central to PAH disease biology that may serve as a therapeutic target as well as a marker of disease prognosis and response to interventions.

The development of advanced tools for defining individuals with PAH at high risk for adverse outcomes is a critically unmet need that is necessary to guide clinical follow-up, accelerate treatment, and facilitate advanced therapies, such as lung transplantation, where timely interventions are resource limited. Moreover, identification of novel biomarkers as more precise and early surrogate endpoints for mortality or transplantation could be utilized in clinical trials to advance precision medicine approaches in this disease. Current attempts to stratify PAH patients rely predominantly on clinical and hemodynamic parameters with several clinical risk scores and calculators to classify patients at low, intermediate, or high risk of death.(*32, 35, 36*) While widely used in the clinical management of PAH, these contemporary risk predictors have limitations. Greater than 70% of PAH patients are classified into intermediate risk groups using currently available clinical risk scores,(*34*) while our results suggest the utility of more precisely stratifying intermediate clinical risk patients using a PAH-PMRS. When compared to current risk scores, PAH-PMRS correctly reclassified 48% of intermediate risk patients who ultimately died as high-risk, suggesting clinical utility of PAH-PMRS over clinical risk scores. Importantly, the PAH-PMRS was found to replicate across both large, multi-center studies as well as independent, single-center studies, and across subgroups of PAH, from treatment naïve patients to those requiring IV therapy. Given the cross-sectional nature of our study, there is limited ability, however, to detect temporal associations between PPOBAS markers and response to therapy or change in clinical status. Future longitudinal studies are warranted to explore the dynamic nature of these metabolites in relation to PAH progression and treatment response. In particular, recent advancements in pharmacologic modulation and diagnostic assessment of the BMP/TGF-β pathways are poised to offer encouraging clinical improvements to therapy and management,(*37, 38*) and addition of the PAH-PMRS to emerging BMP/TGF-β biomarkers may provide for even further prognostic insight.

Consistent with our findings in PAH, there is growing evidence that primary oxysterol and bile acid pathways are central to fundamental physiologic and pathophysiologic processes, including those external to the liver. In addition to a known role in lipid absorption, oxysterols, bile acids, and their intermediates function as signaling molecules that regulate energy metabolism, inflammation, and immune response—all key molecular determinants of PAH.(*39*) Importantly, primary bile acid synthesis via oxysterol intermediates represents the major catabolic mechanism for cholesterol trafficking. While traditionally centralized to the liver, such processes can occur extrahepatically, such as the lungs, through the activity of expressed CYP7B1 and/or CYP27 enzymes.(*40–42*) Furthermore, there is an increasing appreciation of the proinflammatory and pathogenic actions of oxysterols and bile acids in vascular diseases.(*43–46*) In the context of PAH, prior evidence points to dysregulated cholesterol transport and processing. Namely, patients with PAH have been reported to display lower circulating LDL cholesterol but increased circulating and lung tissue oxidized LDL and oxidized LDL/LDL ratio.(*47*) In fact, PAH lung tissues carry reduced levels of the LDL receptor (LDLR), a molecule that controls the initiating step for cholesterol internalization, transport to the lysosome, and downstream metabolism. In a small study on human lung tissues, PAH patients displayed higher expression of CYP7B1—the initial and rate-determining enzyme in the classic pathway of bile acid synthesis—and elevated levels of bile acids metabolites when compared to heathy controls,(*40*) supporting the findings herein. Most importantly, leveraging the findings of our multi-omic analyses described here, our accompanying study (by Harvey *et al.*)(*20*) defines a causal role for PPOBAS within the endothelium as a crucial inflammatory driver of PAH severity and a pathway that can be pharmacologically targeted for therapeutic benefit. Taken together, such findings offer convergent evidence of the mechanistic underpinnings linking PPOBAS with PAH pathobiology that can be leveraged for development of precision medicine approaches in diagnosis and therapy.

Several limitations of the study merit consideration. The cross-sectional design of our study limits the ability to detect temporal associations between the metabolites and response to therapy or change in clinical status. Although enrollment carried representation across 28 institutions with diverse ethnic and racial representation, the extent to which findings may be generalized beyond the United States is unclear. The untargeted nature of our metabolomics analysis limits identification of all measured metabolites, suggesting that additional analytes may be found in future work to carry robust associations with either risk or severity of PAH. Finally, while this study focuses on defining a prognostic PAH-PMRS score driven by a metabolome-wide association of key metabolites with PAH severity, our companion study more specifically defines the mechanistic underpinnings linking these oxysterols and bile acid intermediates with PAH severity and the utility of therapeutic targeting. Our PAH cohorts included patients with already diagnosed and clinically phenotyped PAH, and so additional studies are needed to evaluate the extent to which PPOBAS pathway is detectable and clinically informative in earlier subclinical stages of PAH development. Additional follow-up studies are also needed to identify molecular signatures and drivers of risk that may be specific for a given etiologic subtype of PAH, notwithstanding the potential role of PPOBAS in representing a common pathway to morbidity and mortality across PAH subtypes.

In conclusion, our study leverages large-scale, unbiased metabolomic analyses to define PPOBAS intermediates as key prognostic markers and determinants of PAH severity and prognosis. Beyond the identification of the 51 PPOBAS metabolites associated with PAH risk and mortality, our study also revealed numerous novel and not previously annotated metabolites that are significantly associated with PAH mortality, including those that validate across independent cohort studies. Future work is needed to assess the potential of these metabolites to serve therapeutic targets and or measures of treatment response as part of ongoing efforts to develop more effective precision medicine approaches for the care of patients with PAH. More broadly, guided by prior predictions of this platform,(*48*) our work now demonstrates both the feasibility and the expansiveness of large-scale mass spectrometry analysis and multi-dimensional data integration in the application to human disease.

## Acknowledgments

We thank our collaborators, and their Pulmonary Hypertension Centers, who collected samples used in this study, as well as patients and their families, whose help and participation made this work possible. We thank all participants of the FINRISK 2002 survey for their contributions to this work. Schematic study design in Fig. 1. and schematic bile acid pathway in Fig. 2D were created with BioRender.com.

## Data and materials availability

All data, code, and materials used in the analysis are available to researchers upon reasonable request and following issuance of relevant institutional data sharing agreements. In particular, PAH Biobank data are publicly available to bona fide researchers upon application at http://www.pahbiobank.org. The FINRISK data for the present study are available with a written application to the THL Biobank as instructed on the website of the Biobank (https://thl.fi/en/web/thl-biobank/for-researchers). A separate permission is needed from FINDATA (https://www.findata.fi/en/) for use of the EHR data.

