## supplemental methods for "Pulmonary primary oxysterol and bile acid synthesis as a predictor of outcomes in pulmonary arterial hypertension"

***Study cohorts, sample and data acquisition.***

This is a multicenter, observational cohort study. Comprehensive, non-targeted metabolomics analysis of plasma samples from five cohorts of patients with PAH and controls was performed. Cohorts 1 and 2 were enrolled as part of the US National Biological Sample and Data Repository for Pulmonary Arterial Hypertension/PAH Biobank (PAHB) study. The PAH Biobank is an ongoing NIH-funded (R24HL105333, R01HL160941) biorepository of PAH patients enrolled from 37 US centers with deidentified clinical data, banked biological samples and genetic data. In total, 2900 men and women were enrolled between October 2012 to December 2017. A total of 2490 patients with PAH met the inclusion criteria and had baseline plasma biospecimen and were included in this study. Non-fasting blood samples were collected at the time of enrollment. Patients were divided into discovery (cohort 1) and validation 1 (cohort 2) cohorts, *a priori,* based on centers of origins. The discovery cohort included 1,913 patients and the validation cohort included 575 patients. Cohort 3 was enrolled from Vanderbilt Pulmonary Hypertension Research Cohort.(*1*) 102 patients with idiopathic or heritable PAH were enrolled between July 2007 to October 2012. Cohort 4 was enrolled from the University of Pittsburgh Medical Center (UPMC) pulmonary arterial hypertension center. 122 patients were enrolled between April 2016 to October 2018. 44 healthy and disease controls were recruited from UPMC. Disease controls were patients presented with symptomatic dyspnea and exercise intolerance and later confirmed not to have pulmonary hypertension. Diagnosis of PAH was defined by right heart catheterization measurements including mPAP > 25 mmHg, PCWP ≤ 15 mmHg, and PVR > 3 Woods Units, based on the prevailing diagnostic criteria for PH at the time of cohort recruitment.(*2*) Combining cases with controls data, a total of 2,756 subjects were available for analysis. The FINRISK-2002 cohort represents an independent, prospective population survey within the Finnish National FINRISK study (Coordinating Ethical Committee of the Helsinki and Uusimaa Hospital District, Ref. 558/E3/2001). Blood samples were obtained from 8,738 individuals (54% females) with baseline age between 24 to 75 years. Genotyping was performed on Illumina genome-wide SNP arrays (the HumanCoreExome BeadChip, the Human610-Quad BeadChip and the HumanOmniExpress) as described previously.(*3-5*) All studies were conducted in accordance with the Declaration of Helsinki, V edition (2000). The study protocol was approved by the institutional review boards of all the participating centers and all study participants signed informed consent using the PAHB consent forms (R24HL105333), or local institutional consent forms.

***Animal studies.***

All animal experiments were approved by the University of Pittsburgh. To induce pulmonary hypertension (PAH), monocrotaline and Sugen/hypoxia PAH models were used as previously described.(*6*)

*Monocrotaline-PAH model:* Male Sprague Dawley rats (10–14 weeks old, Charles River) were injected i.p. with 60 mg/kg monocrotaline (Sigma-Aldrich) at time 0. On day 21 after monocrotaline injection, right heart catheterization was performed followed by harvesting of lung tissue and obtaining plasma samples.

*Sugen/hypoxia PAH-model:* Male Sprague-Dawley rats (10-14 week old, Charles River) were injected with SU5416 (20mg/kg; Sigma-Aldrich), followed by exposure to normobaric hypoxia (10% O2;OxyCycler chamber, Biospherix Ltd, Redfield, NY) for 3 weeks. Right heart catheterization was performed, followed by harvest of lung tissue and obtaining plasma samples.

***PAEC studies.***

Primary human (Lonza, #CC-2530) pulmonary arterial endothelial cells (PAECs) were grown in EBM-2 basal medium supplemented with EGM-2 MV BulletKit (Lonza). Experiments were performed at passages 5 to 8. Silencer siRNAs for BMPR2 (AM16708) and scrambled control (12935300) were purchased from Thermo Fisher Scientific. PAECs were plated in collagencoated plastic and transfected 16h later at 70-80% confluence using siRNA (20nM) and Lipofectamine 2000 reagent (Thermo Fisher Scientific), according to the manufacturer's instructions. 48 hours post-transfection, cells were lysed for metabolomic analysis.

***Sample preparation.***

Plasma samples were thawed at 4C overnight; 20 µL of plasma was transferred into a 96-well extraction plate. Proteins were precipitated with the addition of 80 µL of ice-cold extraction, as described.(*7*) Plates were sealed, vortexed and centrifuged to precipitate proteins with the supernatant containing extract metabolites pipetted into a clean microtiter plate for liquid chromatography-mass spectrometry (LC-MS) analysis. Samples were passed across a reverse phase SPE column (Phenomenex 8B-S199-UB), and lipid species eluted using 1ml methanol, as described,(*7*) before transferred into a clean microtiter plate for LC-MS analysis. Pre-weighed frozen tissue samples were homogenized using an Omni International Bead Ruptor Elite homogenizer with 300 mg of 1 mm zirconium disruption beads. Ethanol:water (80:20) extraction solvent, containing deuterated internal standards, was added at volumes to achieve a normalized concentration of 50 mg of tissue per 1 µL of solvent for each sample. The homogenate was then transferred into an Axygen 500 µL V-bottom 96-well plate (P/N P-DW-500-C) with 300 µL of water in each well. Samples were passed across a reverse phase SPE wells and lipid species eluted using methanol, as described, before transferred into a clean microliter plate for LC-MS analysis. Cell samples were centrifuged at 14,000 g at 4°C for 10min then the supernatant was transferred to microcentrifuge tube. Samples were resuspended in 80:20 methanol:water and transferred to LC-MS vials containing a 200 µL glass inserts, as described previously.(*8*) Media samples were prepared by taking 50 µL of media and adding 200 µL of ice cold methanol, centrifuged and supernatant was then transferred to LCMS vials containing 200 µL glass inserts, as described.(*8*)

***Metabolite assay.***

Bioactive metabolites analysis was performed on plasma, tissue, cell and media samples using state of the art liquid chromatography-mass spectrometry (LC-MS), using a Vanquish UPLC coupled to high resolution, QExactive orbitrap mass spectrometer (Thermo) using a Phenomenex Kinetex C18 column. All generated spectral data underwent daily Qc/Qa analysis, as described in the section below. Data was extracted using image processing and machine learning based spectral optimization. Each sample MS data file was initially represented as an image using mass to charge and retention time coordinates. A watershed algorithm was applied to image files using peak apex density as a guide to define regions containing putative spectral peaks. An optimized neural network was subsequently applied to all putative spectra peaks to isolate true spectral peaks from ‘false’ background signals. Following data extraction, spectral peaks were cross aligned among datasets using landmark based algorithms to allow for comparison of signals among cohorts. Data was subsequently normalized to account for plate-to-plate variation using a simple batch median normalization metric with correction for median absolute deviation. Following normalization, metabolite peaks were further compressed for multiple adducts and in source fragments. Normalized, aligned, filtered datasets were subsequently used for statistical analyses, as described below. Metabolites were annotated by using an in-house library of commercially available standards or MS/MS fragmentation patterns.

***QC/QA of spectral data.***

Qc/Qa of data has been standardized using a panel of deuterated internal standards as well as interval pooled plasma samples to monitor fluctuations in extraction efficiency, instrument sensitivity, matrix artifact and mass accuracy. Any samples not meeting Qc/Qa thresholds underwent reinjection. The assay was highly reproducible over several days and independent measures with median coefficient of variance (CV) across analytes at low standard concentrations of 0.15 ng.

***Metabolites identification.***

Metabolites meeting strict thresholds for association with clinical phenotypes, as described below, were subsequently evaluated for structural elucidation. Peak identity and isobaric purity were confirmed by comparing MS/MS fragmentation data as well as directionality and intensity correlations with phenotypes. For metabolites meeting this initial threshold, targeted MS/MS fragmentation was preformed and first searched against our internal library of >2000 commercial standards as well as searched against public spectral metabolite libraries, including Metlin (http://metlin.scrips.edu), Massbank (http://www.massbank.jp) and HMDB (http://www.hmdb.ca). Remaining unidentified signals were further characterized by manual MS/MS deconstruction analysis from $MS^n$ multi-level fragmentation data. Commercial or synthesized standards were purchased for putative biomarkers. Microsomal glucuronidation was performed on 25 oxysterols and bile acid intermediates standards and matched with selected metabolites using exact mass to charge (M/Z), retention time (RT) and peak shape.

***Molecular networking.***

Spectral networking was employed to subclassify metabolites of interest into those sharing similar chemical structure.(*9-11*) This approach ‘blasts’ MS/MS spectral patterns obtained from each small molecule against a database of over 1 billion experimentally obtained spectra, matching each small molecule according to chemical similarity to ‘reference knowns’ and thereby providing clues into the origins of each plasma small molecule.  Vector similarities was calculated for every possible pair of MS/MS spectra between small molecules of interest and the MassIVE GNPS database, containing MS/MS spectra from over 40,000 pure standards and 1.2 billion user obtained MS/MS spectra (http://gnps.ucsd.edu). For molecular networking, cosine threshold values were set at 0.6-0.7 where a cosine value of 1.0 indicates identical spectra. Similarity scores was visualized by plotting them on a 2D plane using Cytoscape.(*12*)

***Statistical analysis.***

Continuous variables of demographic data are reported as mean +/- SD. The primary endpoint of the study was composite endpoint of all-cause mortality or lung and heart transplantation. Survival time was defined as the time from sample collection to the date of death or transplantation, censored at the last follow-up or alive until the last follow-up. Prior to all analyses, metabolite values were natural logarithmically transformed, as needed, and later standardized with mean=0 and SD=1 to facilitate comparisons. Cox regression analysis was used to examine the association between each metabolite level (independent variable) and the primary outcome (dependent variable, all-cause mortality or lung and heart transplantation). Two sets of covariates were used. The first set included age, sex and BMI. The second set included age, sex, BMI, traditional clinical risk factors such as cirrhosis, renal failure, and medication use including prostacyclin, aspirin, and statin. Correction for multiple testing was performed using the Bonferroni-Holm method. Benjamin-Hochberg false discovery rate (FDR) was also calculated and metabolites not meeting threshold of $q < 0.01$ were excluded. Metabolites meeting significance threshold were evaluated in the validation cohort (cohort 2) by Cox regression analysis, adjusting for the previous covariates. In the validation cohort, the significance threshold was recalculated based on the significant metabolites from the discovery cohort (0.05 divided by the number of significant metabolites from the discovery cohort, 1E-4) with consistent direction of effect size. Linear additive models were performed between the selected metabolites and previously reported single nucleotide polymorphism (SNP) genotyping in the FINRISK-2002 cohort. Specifically, SNPs were previously determined on a genome-wide scale for this population, using Illumina genome-wide SNP arrays (the HumanCoreExome BeadChip, the Human610-Quad BeadChip and the HumanOmniExpress). (*4, 5*)Linear and logistic regression analyses were performed between the selected metabolites and clinical markers of PAH severity including 6-minute walk distance (6MWD), WHO functional class (FC), mean right atrial pressure (mRAP), pulmonary vascular resistance (PVR) and cardiac index (CI). To construct a prediction model based on the minimum number of metabolites meeting the statistical threshold, regularized regression was implemented using the LASSO model with 10-fold cross validation. The penalty parameter was chosen as the $\lambda$ that gave the minimum error as determined by the 10-fold cross validation. Receiver operating characteristic (ROC) curves were constructed, and C statistics were calculated using the 13 selected metabolites, clinical covariates alone (including age, gender, BMI, mPVR, mRAP, CI, WHO FC) and metabolites plus clinical covariates. The discovery cohort was used as the training set, cohort 2 as the first testing set and cohort 3 as the second testing set. Metabolites selected by LASSO method were used to construct the PAH-poly metabolite risk score (PAH-PMRS). PAH-PMRS was calculated based on the formula: $\beta_1X_1+\beta_2X_2+...+\beta_nX_n$ with Xn denoting the standardized value for the selected metabolites abundance, and $\beta_n$ denoting the log transformed regression coefficient from the regression model for the primary outcome containing the statistically significant metabolites.(*13, 14*) The scores were grouped into tertiles, and tested using a class variable to estimate the odds ratio for each tertile. Patients having risk scores smaller than the lower tertile were considered low risk group, patients having risk scores larger than top tertile were considered high risk group and all the rest were considered medium risk group. Kaplan-Meier estimates were produced for strata based on PAH-PMRS. Log-rank test was used to compare survival analysis between the groups. Statistical analysis was performed with R with RStudio and associated packages.

**Suppl Fig S1. Spectral network analysis of PAH mortality associated metabolites.**

Expanded view of the largest network from (**Fig. 1B**) showing chemical clustering with glucuronide molecules. Overlay of tandem mass spectra from sterol-glucuronide molecule and m/z 611.3360 showing a high degree of similarity.

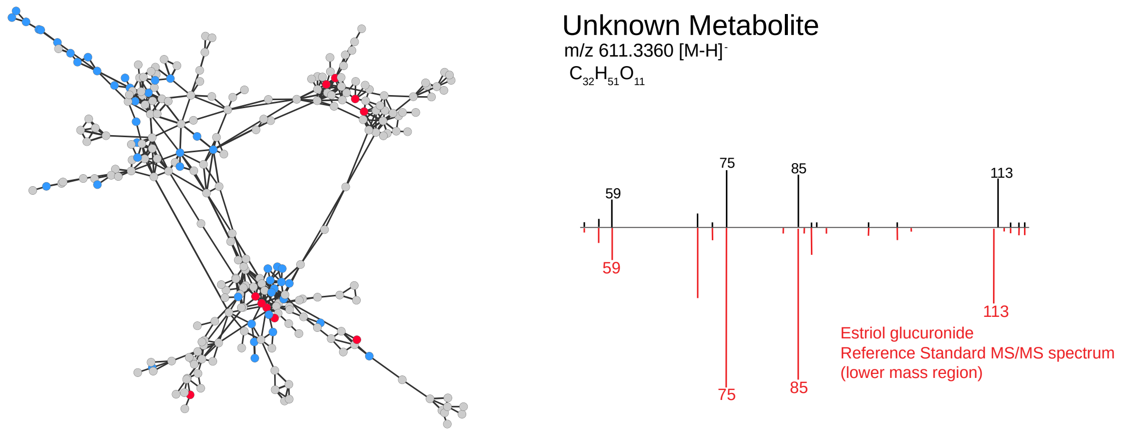

**Suppl Fig S2. Heatmap of Pearson's correlations among 51 glucuronidated metabolites.**

Correlation matrix and cluster analysis of 51 glucuronidated metabolites. Each square indicates a Pearson's correlation coefficient for a pair of metabolites. The value for the correlation coefficient is represented by the intensity of the blue or red color, as indicated on the color scale. Hierarchical clusters are presented as a cluster tree.

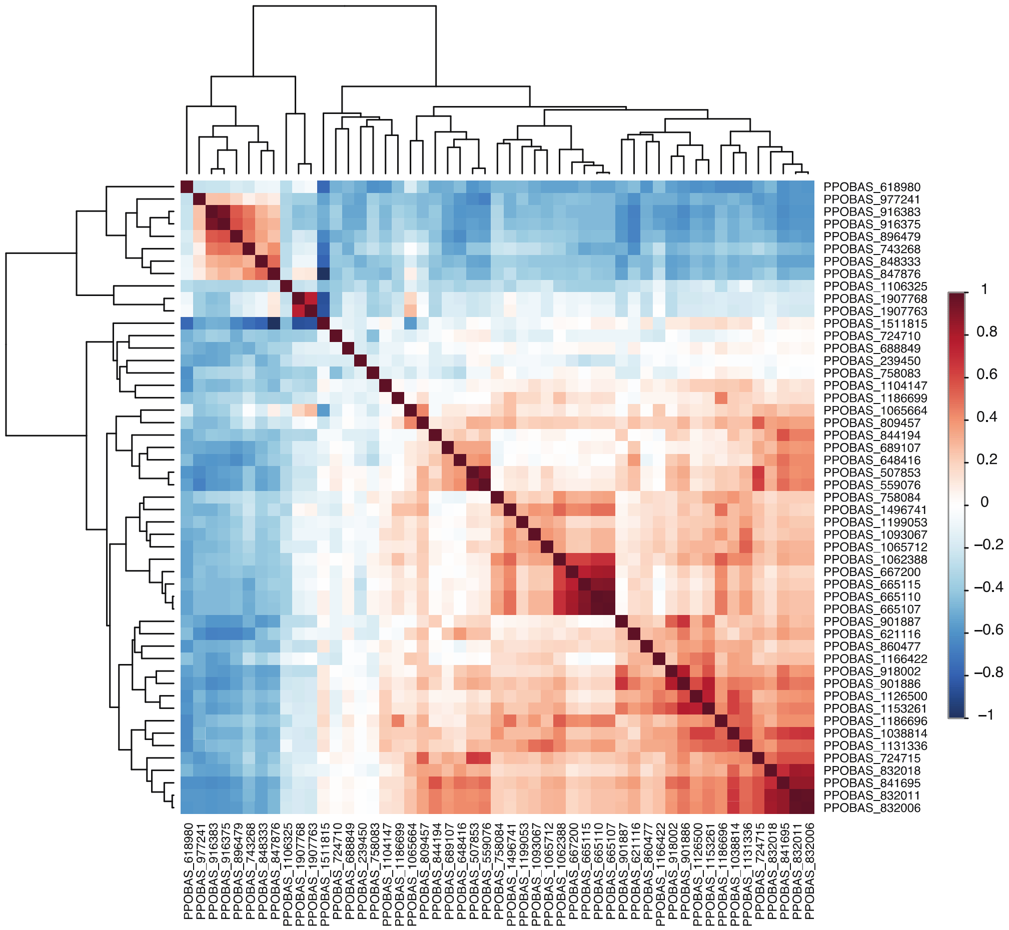

**Suppl Fig S3. Reclassification of PAH clinical risk score using PAH-PMRS in an independent validation cohort.**

Reclassification of PAH clinical risk score using European Society of Cardiology/European Respiratory Society (ESC/ERS) PAH risk calculator in patients who died in cohort 3 (N=27/102) using PAH polymetabolite risk score (PAH-PMRS).

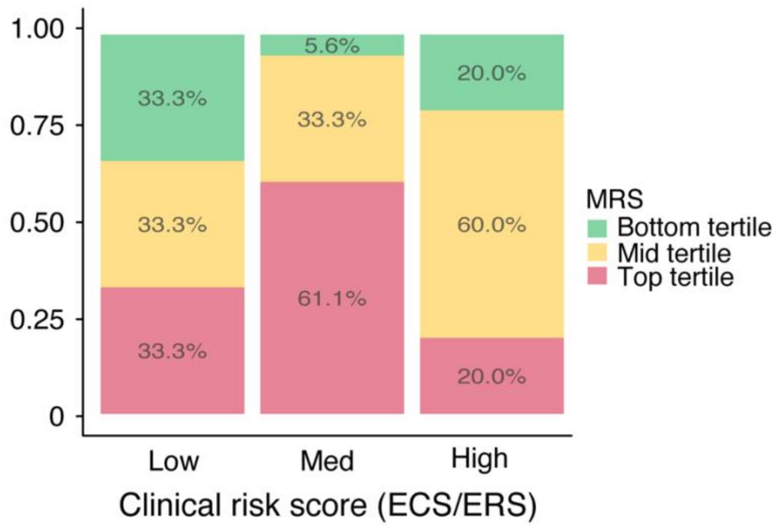
